## Supplementary Legends for "Mutant mice lacking alternatively spliced p53 isoforms unveil *Ackr4* as a male-specific prognostic factor in Myc-driven B-cell lymphomas"

**Supplemental Information**

**Legends to Supplementary Figures and Tables**

**Figure S1.** **Description of the p53^ΔAS^ mouse model and analysis of p53^ΔAS/ΔAS^ thymocytes and fibroblasts.**

(**A**) Relative expression of p53 isoforms with an α or AS C-terminus in tissues of wild-type mice. RNAs were prepared from the indicated tissues of wild-type mice, then p53-AS and p53-α mRNAs were quantified by RT-qPCR. Results are expressed as mean AS/α ratios ± SD, from 3 independent experiments. (**B**) Comparative maps of the wild-type (WT) and ΔAS *Trp53* alleles. Left, top: the 3’ end of the WT *Trp53* gene is shown - black line: intron 10; boxes are for exons, with greytones for translated regions from exon 10 (light gray), exon 11 (dark gray), and exon AS (black). Left, below: the WT *Trp53* allele encodes proteins with two different C-termini. (4D: tetramerization domain, CTD: C-terminal domain). Right, top: the 3’ end of the ΔAS *Trp53* allele is shown; the AS exon was deleted and replaced by a SpeI restriction site. Right, below: the ΔAS *Trp53* allele only enables the synthesis of proteins with the ‘canonical’ α C-terminus. (**C**) Quantifications of p53 isoforms with an AS or an α C-terminus, in mouse embryonic fibroblasts (MEFs) of the indicated genotypes, that were either untreated, or treated for 24h with 0.5 μg/ml doxorubicin (Doxo) or 10 μM Nutlin. Means ± SEM from >3 experiments with ≥ 2 independent MEF clones of each genotype are shown. ***P<0.001, *P<0.05, °P=0.07 by Student’s t-test. (**D**) Apoptotic responses of p53^+/+^ and p53^ΔAS/ΔAS^ thymocytes. p53^+/+^ and p53^ΔAS/ΔAS^ mice were irradiated and their thymocytes were recovered and analyzed by FACS after annexin V-FITC staining. A typical experiment for cells of each genotype and condition is shown. Numbers indicate % cells, apopt.: apoptotic cells. (**E**) Cell cycle control of p53^-/-^, p53^+/+^ and p53^ΔAS/ΔAS^ fibroblasts. Asynchronous MEFs were exposed to 0-10 Gy γ-irradiation, then after 24 hr cells were labelled with BrdU for 1 hr and analyzed by FACS. A typical experiment for cells of each genotype and condition is shown, with % of cells in G1, S and G2/M mentioned in each panel. (**F**) mRNA levels of the indicated genes were quantified in MEFs of the indicated genotypes left untreated or treated for 24h with 0.5 μg/ml Doxorubicin (Doxo), 10 μM Nutlin-3 (Nutlin). Means ± SEM from at least 3 experiments with 2 independent MEF clones of p53^+/+^ and p53^ΔAS/ΔAS^ genotypes are shown. (**G**) mRNA levels of the indicated genes were quantified in MEFs of the indicated genotypes left untreated or treated for 24h with 15 μM etoposide (Eto). Means ± SEM from at least 3 experiments with 2 independent MEF clones per genotype.

**Figure S2. Analysis of tumors from p53^+/+^ Eμ-myc and p53^ΔAS/ΔAS^ Eμ-Myc mice.**

(**A**) Histological analyses of Eμ-Myc-induced tumor lymph nodes. Tumor lymph nodes were analyzed by Hematoxilin-Eosin staining (H & E), or antibodies against B220 (a B-cell specific marker) or CD3 (a T-cell specific marker). p53^+/+^ Eμ-Myc and p53^ΔAS/ΔAS^ Eμ-Myc mice developed similar B-cell lymphomas, characterized by massive tissue homogenization of the lymph node by B-cells. Scale bars=50 μM. (**B**) Tumor-free survival of p53^+/+^ Eμ-myc and p53^ΔAS/ΔAS^ Eμ-Myc mice are similar when sexes are not considered (n=cohort size). (**C**) Increased tumor-free survival of p53^+/+^ Eμ-myc males compared to p53^ΔAS/ΔAS^ Eμ-Myc males, p53^+/+^ Eμ-Myc females and p53^ΔAS/ΔAS^ Eμ-Myc females (n=cohort size). Statistical analysis with Mantel-Cox test. (**D**) *Trp53* DNA sequencing of p53^+/+^ Eμ-myc and p53^ΔAS/ΔAS^ Eμ-Myc tumors. The portion of the *Trp53* gene encoding the DNA Binding domain (DBD), most frequently mutated in cancers, was sequenced for 14 tumors from p53^+/+^ (WT) Eμ-myc males and 18 tumors from p53^ΔAS/ΔAS^ (ΔAS) Eμ-myc males, and *Trp53* mutations were found in 3/14 and 1/18 tumors, respectively. (**E**) Details on the 4 p53 mutations identified in (D).

**Figure S3.** **Analysis of pre-tumoral spleens.**

(**A**) Cell sorting of B-cell subpopulations by FACS. Splenic cells were incubated with DAPI and the following antibodies: APC rat anti-mouse CD45R/B220, FITC rat anti-mouse CD43, PE rat anti-mouse IgM and BV605 rat anti-mouse IgD. First, the B220+CD43- cells were selected from DAPI negative living cells, subsequently yielding 4 different B subpopulations based on IgM and IgD labeling: IgM-/IgD- pre-B cells (box 1), IgM low/IgD- immature B cells (box 2), IgM high/IgD- transitional B cells (box 3) and IgM+/IgD+ mature B cells (box 4). Typical results with a p53^+/+^ Eμ-Myc male mouse and a p53^ΔAS/ΔAS^ Eμ-Myc male mouse are shown. (**B**) The loss of p53-AS isoforms does not impair B-cell differentiation in 6 weeks-old non-transgenic male mice. Means ± SEM from 4 mice per genotype. ns: non-significant by Student’s t-test. (**C**) Control for expression of p53-FL, p53-AS and p53^R270H^ in luciferase assays. As a control to luciferase experiments reported in Fig. 3H, protein extracts from p53^-/-^ MEFs transfected with expression plasmids for p53-FL, p53-AS or p53^R270H^ were immunoblotted with antibodies against p53, p21 and actin. Quantifications are relative to actin. p53-FL and p53-AS were expressed at similar levels and both transactivated p21, whereas the DNA-binding mutant p53^R270H^ was expressed at higher amounts but failed to transactivate p21, as expected. nd: not determined.

**Figure S4. Further analyses of human B-cell lymphoma or Multiple myeloma datasets.**

(**A**) *MT2A* gene expression is not a prognostic marker in human B-cell lymphomas. Survival curves for the 30% patients with the highest *MT2A* mRNA levels and the 30% patients with the lowest *MT2A* mRNA levels according to sex (n=cohort sizes), for patients from dataset #GSE4475. (**B**) *ACKR4* is a prognostic factor in Burkitt lymphomas but not in diffuse large B-cell lymphomas. Survival curves of patients from dataset #GSE181063, for the 30% patients with the highest *ACKR4* mRNA levels and the 30% patients with the lowest *ACKR4* mRNA levels, classified according to sex (n=cohort sizes), and diagnosed with either a diffuse large B-cell (top) or a Burkitt (bottom) lymphoma. (**C**) *ACKR4* is not a prognostic factor in Multiple Myeloma. Survival curves of patients from dataset #GSE136337, for the 30% patients with the highest *ACKR4* mRNA levels and the 30% patients with the lowest *ACKR4* mRNA levels in malignant plasma cells, classified according to sex (n=cohort sizes). Statistical analyses in all panels by Mantel-Cox tests.

**Figure S5. Strategy to knockout *ACKR4* in Burkitt lymphoma cells.**

Burkitt lymphoma cells were transfected with a PX459 vector expressing Cas9, a puromycin resistance gene and either of two guide RNAs targeting *ACKR4* (or no guide RNA for control), then puromycin-resistant cells were selected and recovered either as cellular pools or diluted to get 1 cell per 5 wells in a 96-well plate to isolate cellular clones. Individual clones were then expanded, and clonal cell populations were split for freezing and DNA extraction. DNA was analyzed by performing a PCR amplifying a fragment of *ACKR4*, then amplified products were cloned in a PGL3 plasmid for DNA sequencing. For each cellular clone, 8 plasmid clones were sequenced to ensure information on both *ACKR4* alleles.

**Figure S6. Characterization of *ACKR4* KO Burkitt lymphoma cell clones.**

(**A**) Characterization of the *ACKR4* KO 4.14 clone. Left, DNA sequences of the region targeted by the guide RNA g4. Compared to the WT *ACKR4* DNA sequence from Burkitt lymphoma (BL) Raji control cells (center), the 4.14 clone has two mutated alleles: allele a, with a deletion of 17 nt (top) and allele b, with a deletion of 4 nt (bottom). Right, comparison of the ACKR4 proteins encoded by WT alleles from BL Raji control cells (center), or by the 4.14a (top) and 4.14b mutated alleles (bottom). The WT ACKR4 protein consists of 350 amino acids, including a DRY motif (at residues 136-138) essential for signal transduction. The protein region corresponding to the target of guide RNA g4 is indicated. Allele 4.14a encodes a putative protein with 59 N-terminal residues of ACKR4 and 34 amino acids of unrelated sequence due to the mutational frameshift. Allele 4.14b encodes a putative protein with 60 N-terminal residues of ACKR4. (**B**) Characterization of the *ACKR4* KO 5.2 clone. DNA sequences of the region targeted by the guide RNA g5 (left) and comparison of the encoded ACKR4 proteins (right), are represented as in (A).

**Figure S7. The knockout of *ACKR4* in Burkitt lymphoma Raji cells does not impact their proliferation.**

Equal numbers of cells of the indicated genotypes were seeded, then cultured for 15 h with or without CCL21 and counted. Statistical analyses by Student’s t test. ns: non-significant.

**Table S1. Expression of *Ackr4*, *Cdkn1a* and *Mdm2* in p53^+/+^ Eμ-Myc and p53^ΔAS/ΔAS^ Eμ-Myc male splenic cells.**

Read numbers for the indicated genes, obtained by Bulk RNA-seq from the spleens of three p53^+/+^ Eμ-Myc (WT_Myc) and four p53^ΔAS/ΔAS^ Eμ-Myc (ΔAS_Myc) male mice.

**Table S2. Details of the gene set enrichment analysis (GSEA) for hallmark Myc targets.**

Datasets from splenic cells of three p53^+/+^ Eμ-Myc and four p53^ΔAS/ΔAS^ Eμ-Myc male mice were analyzed by GSEA. A normalized enrichment score of 2.4965038 (with a false discovery rate of 0.0) was found for the gene set ‘Hallmark_Myc_targets_V1’ in p53^ΔAS/ΔAS^ Eμ-Myc cells. The table provides details on the profile represented in Figure 2K, with running enrichment scores and positions of gene set members on the rank ordered list.

**Table S3.** **Oligonucleotide Sequences.**
