## Supplemental Table S1 for "Mutant mice lacking alternatively spliced p53 isoforms unveil *Ackr4* as a male-specific prognostic factor in Myc-driven B-cell lymphomas"

| **Gene** | **WT_Myc_1** | **WT_Myc_2** | **WT_Myc_3** | **ΔAS_Myc_1** | **ΔAS_Myc_2** | **ΔAS Myc_3** | **ΔAS _Myc_4** |
| --- | --- | --- | --- | --- | --- | --- | --- |
| *Ackr4* | 267.610698766789 | 187.913317967838 | 103.698438238889 | 80.2839670298248 | 58.3264519788058 | 44.2019721676723 | 58.626719832054 |
| *Cdkn1a* | 1172.3897279307 | 1179.72136964871 | 1227.565295909 | 1248.51861547663 | 1018.37985154995 | 800.288338193645 | 1079.90417930643 |
| *Mdm2* | 4419.00743604647 | 4820.24950903391 | 5133.53980290714 | 4756.31040570283 | 4240.33305885918 | 4132.88439767736 | 5521.46447378284 |

**Table S1. Expression of *Ackr4*, *Cdkn1a* and *Mdm2* in p53^+/+^ Eμ-Myc and p53^ΔAS/ΔAS^ Eμ-Myc male splenic cells.**

Read numbers for the indicated genes, obtained by Bulk RNA-seq from the spleens of three p53^+/+^ Eμ-Myc (WT_Myc) and four p53^ΔAS/ΔAS^ Eμ-Myc (ΔAS_Myc) male mice.
