## Supplemental Table S3 for "Mutant mice lacking alternatively spliced p53 isoforms unveil *Ackr4* as a male-specific prognostic factor in Myc-driven B-cell lymphomas"

| Locus | Species | Usage | Forward | Reverse |
| --- | --- | --- | --- | --- |
| *Trp53* | Mouse | Genotyping | 5’ AAGGGGTATGAGGGACAAGG 3’ | 5’ GAAGACAGAAAAGGGGAGGG 3’ |
| *Eμ-Myc* | Mouse | Genotyping | 5’ ACCCAGGCTAAGAAGGCAAT 3’ | 5’ CGCTCACTCCCTCTGTCTCT 3’ |
| *Trp53* exon4 | Mouse | Sequencing | 5’ CAGAGCAGAAAGGGACTTGG 3’ | 5’ GCTAAAAAGGTTCAGGGCAAA 3’ |
| *Trp53* exon5-6 | Mouse | Sequencing | 5’TGGTGCTTGGACAATGTGTT 3’ | 5’TAGCACTCAGGAGGGTGAGG 3’ |
| *Trp53* exon7-9 | Mouse | Sequencing | 5’TGCCGAACAGGTGGAATATC 3’ | 5’CCTTGGTACCTTGAGGGTGA 3’ |
| *Ackr4* | Mouse | RT-qPCR | 5' GCACCTCTCCCAGCTTAAACA 3' | 5' AATAGTATTCCGCTGACTGGTTCAG 3' |
| *Akr1c19* | Mouse | RT-qPCR | 5' AGTTGCCTACTGTGCTCTTGGAT 3' | 5' GAGAACTGGAGAGCTTGGGTCTA 3' |
| *Bax* | Mouse | RT-qPCR | 5' CGGCGAATTGGAGATGAACT 3' | 5' CCGTGTCCACGTCAGCAA 3' |
| *Cdkn1a* | Mouse | RT-qPCR | 5’ GCAGACCAGCCTGACAGATTTC3’ | 5’ TTCAGGGTTTTCTCTTGCAGAAG 3’ |
| *Cd300lh* | Mouse | RT-qPCR | 5' GAACATTGGCCAGGTGACTCA 3' | 5' AAGAAGGAGATGCTGCTCAACAG 3' |
| *Fam132a* | Mouse | RT-qPCR | 5' GACAAGAAGACTCTGGTGGAACTG 3' | 5' AGGAAGGCGCCCTGAGTAGT 3' |
| *Fas* | Mouse | RT-qPCR | 5’ ATGCACACTCTGCGATGAAG 3’ | 5’ CAGTGTTCACAGCCAGGAGA 3’ |
| *Il5ra* | Mouse | RT-qPCR | 5' CCGCCTGCTTCGTCTTGTTA 3' | 5' CAACCTGGTCCATAGATGACACA 3' |
| *Masp2* | Mouse | RT-qPCR | 5' CCAATGAGAAGCCGTTCACA 3' | 5' GAGACACTCTGCATTCATCCACAT 3' |
| *Mdm2* | Mouse | RT-qPCR | 5’ GTCTACCGAGGGTGCTGCAA 3’ | 5’ AAGCAATGGTTTTGGTCTAACCA 3’ |
| *Mt2* | Mouse | RT-qPCR | 5' ACAATGCAAATGTACTTCCTGC 3' | 5' CACTTCGCACAGCCCACG 3' |
| *Myc* | Mouse | RT-qPCR | 5' CCACCAGCAGCGACTCTGA 3' | 5' TCCACAGACACCACATCAATTTC 3' |
| *Noxa* | Mouse | RT-qPCR | 5' CCTGGGAAGTCGCAAAAGAG 3' | 5' GAGCACACTCGTCCTTCAAGTCT 3' |
| *Prss50* | Mouse | RT-qPCR | 5' CGTGGCCCATTGCTTGA 3' | 5' GCTCCCCGCCCTCACT 3' |
| *Puma* | Mouse | RT-qPCR | 5’ GAGCGGCGGAGACAAGAA 3’ | 5’ GAGTCCCATGAAGAGATTGTACATGA 3’ |
| *Redd1* | Mouse | RT-qPCR | 5' GAGTCCCTGGACAGCAGCAA 3' | 5' CATCCAGGTATGAGGAGTCTTCCT 3' |
| *Sesn2* | Mouse | RT-qPCR | 5' CTTCCGCCACTCAGAGAAGGT 3' | 5' CTTGCATGCGGGCTTCA 3' |
| *Slc26a1* | Mouse | RT-qPCR | 5' GCCACTGCCCTTACTCTGATG 3' | 5' GCCGGAGGATACCCATGAG 3' |
| *Survivin* | Mouse | RT-qPCR | 5' TGTTTTTTCTGCTTTAAGGAATTGG 3' | 5' TCTATGCTCCTCTATCGGGTTGTC 3' |
| *Tcstv3* | Mouse | RT-qPCR | 5' CTCCAGCTGTTGTGGAATAAGTTC 3' | 5' CCATGGATCCCTGAAGGTAAATC 3' |
| *Trp53-α* | Mouse | RT-qPCR | 5’ AAAGGATGCCCATGCTACAGA 3’ | 5’ TCTTGGTCTTCAGGTAGCTGGAG 3’ |
| *Trp53-AS* | Mouse | RT-qPCR | 5’ AAAGGATGCCCATGCTACAGA 3’ | 5’ TGAAGTGATGGGAGCTAGCAGTT 3’ |
| *Ulk1* | Mouse | RT-qPCR | 5' TGTACATGGCTCCTGAGGTCAT 3' | 5' CTCCACAGGTCAGCCTTTCC 3' |
| *Ppia* | Mouse | RT-qPCR control (MEFs) | 5' TCTCCTTCGAGCTGTTTGCA 3' | 5' CAGTGCTCAGAGCTCGAAAGTTT 3' |
| *Rplp0* | Mouse | RT-qPCR control (MEFs) | 5' CGACCTGGAAGTCCAACTAC 3' | 5' ATCTGCTGCATCTGCTTG 3' |
| *Il2rg* | Mouse | RT-qPCR control (spleen) | 5' GGAGCTCCAAGGTCCTCATG 3' | 5' TGTAGAAGTCAGGATCAAATCAGCTT 3' |
| *Polr2a* | Mouse | RT-qPCR control (spleen, thymus) | 5' CTTTGAGGAAACGGTGGATGTC 3' | 5' TCCCTTCATCGGGTCACTCT 3' |
| *Vps4a* | Mouse | RT-qPCR control (thymus) | 5' GACAACGTCAACCCTCCAGAAA 3' | 5' TCTGTGGCTTTTGTCACCAGAT 3' |
| *ACKR4* | Human | Sequencing | 5’ AGTGCTAGATTCAGGCTCACA 3’ | 5’ CAGCCATCCAGACACAGAAA 3’ |
| *ACKR4* | Human | RT-qPCR | 5' ACTGCTCCTCTCTGCCGACTAC 3' | 5' GCCATTCATTTCATTTTCCTCAT 3' |
| *CDKN1A* | Human | RT-qPCR | 5' ACCATGTGGACCTGTCACTGTCTT 3' | 5' AGAAGATGTAGAGCGGGCCTTTGA 3' |
| *PPIA* | Human | RT-qPCR control | 5' CATCTGCACTGCCAAGACTGA 3' | 5' TTCATGCCTTCTTTCACTTTGC 3' |
| *RPLP0* | Human | RT-qPCR control | 5' CTTGTCTGTGGAGACGGATTACAC 3' | 5' TCAGCCAAGAAGGCCTTGA 3' |
| *ACKR4* | Human | *ACKR4* KO (gRNA) *#4* | 5’ TGGTAGTGGCAATTTATGCC 3’ | 5’ GGCATAAATTGCCACTACCA 3’ |
| *ACKR4* | Human | *ACKR4* KO (gRNA) *#5* | 5’ GGGCTGTTAATGCAGTTCAT 3’ | 5’ ATGAACTGCATTAACAGCCC 3’ |

**Table S3.** **Oligonucleotide Sequences.**
